# Multiple evolutionary pathways lead to vancomycin resistance in *Clostridioides difficile*

**DOI:** 10.1101/2023.09.15.557922

**Authors:** Jessica E. Buddle, Rosanna C. T. Wright, Claire E. Turner, Roy R. Chaudhuri, Michael A. Brockhurst, Robert P. Fagan

**Affiliations:** Molecular Microbiology, School of Biosciences, University of Sheffield, Sheffield S10 2TN, UK; Division of Evolution and Genomic Sciences, University of Manchester, Manchester M13 9PT, UK

## Abstract

*Clostridioides difficile* is an important human pathogen, for which there are very limited treatment options, primarily the glycopeptide antibiotic vancomycin. In recent years vancomycin resistance has emerged as a serious problem in several Gram positive pathogens, but high level resistance has yet to be reported for *C. difficile*, although it is not known if this is due to constraints upon resistance evolution in this species. Here we show that resistance to vancomycin can evolve rapidly under ramping selection but is accompanied by severe fitness costs and pleiotropic trade-offs, including sporulation defects that would be expected to severely impact transmission. We identified two distinct pathways to resistance, both of which are predicted to result in changes to the muropeptide terminal D-Ala-D-Ala that is the primary target of vancomycin. One of these pathways involves a previously uncharacterised D,D-carboxypeptidase, expression of which is controlled by a dedicated two-component signal transduction system. Our findings suggest that while *C. difficile* is capable of evolving high-level vancomycin resistance, this outcome may be limited clinically due to pleiotropic effects on key pathogenicity trains. Moreover, our data provide a mutational roadmap to inform genomic surveillance.

## Introduction

*Clostridioides difficile* is the most common cause of antibiotic-associated diarrhoea worldwide, resulting in significant morbidity and mortality^1^ that places a huge burden on healthcare systems^2,3^. Most cases of nosocomial *C. difficile* infection (CDI) follow recent antibiotic treatment, which, through altering microbial diversity in the colon, reduces microbiota-mediated colonisation resistance^4^. Although restoring microbial community diversity through faecal microbiota transplantation (FMT) is a potential future treatment for CDI^5^, current treatment relies on additional antibiotics, most commonly metronidazole or vancomycin. While these can resolve the CDI, they further exacerbate damage to the microbiota, leading to recurrent CDI in up to 25% of cases^6^. Due to increasing incidence of resistance and consequently poor patient outcomes, use of metronidazole has declined in recent years, and vancomycin is now the recommended front line antibiotic in the UK^7^.

Vancomycin is a glycopeptide antibiotic that binds to the terminal D-Ala-D-Ala residues on peptidoglycan muropeptide precursors, sterically blocking transglycosylation and transpeptidation reactions^8^. Although relatively slow to emerge, resistance to vancomycin is now found globally in *Staphylococcus aureus* (VRSA) and is widespread in several *Enterococcus* spp. (VRE)^9,10^. Vancomycin resistance in Enterococci is usually associated with one of a number of *van* gene clusters which encode the enzymes that modify peptidoglycan to remove the vancomycin binding site. Here, the D-Ala-D-Ala is replaced with either D-Ala-D-Lac (e.g. *vanA*, also seen in VRSA), conferring high level resistance, or D-Ala-D-Ser (e.g. *vanG*), conferring low level resistance^11^ Vancomycin susceptibility of *C. difficile* is not routinely tested in clinical laboratories making monitoring the emergence of resistance extremely challenging. However, the effectiveness of vancomycin against CDI has declined over time^12^ and multiple case reports show reduced vancomycin susceptibility in individual clinical isolates^13^. Although a complete *vanG* cluster is found in diverse *C. difficile* strains^14^, whether this confers vancomycin resistance is unclear^15^. However, increased expression of the *vanG* cluster does appear to be associated with reduced susceptibility to vancomycin. Moreover, mutations in the *vanSR*-encoded two component system that reduce vancomycin susceptibility through derepression of the *vanG* cluster, have been reported in both clinical isolates and laboratory evolution experiments^16,17^. Beyond such regulatory changes of the *vanG* cluster, the molecular mechanisms underpinning the evolution of vancomycin resistance in *C. difficile* remain unknown. For example, we do not know if other mutations occurring within the *vanG* cluster or at other loci in the *C. difficile* genome contribute to increasing resistance observed clinically. Moreover, whether the evolution of vancomycin resistance is associated with pleiotropic phenotypic effects or fitness costs in *C. difficile* is poorly understood.

To understand the evolutionary dynamics and molecular mechanisms of vancomycin resistance in *C. difficile* we experimentally evolved ten replicate populations at increasing concentrations of vancomycin. Within just 250 generations we observed the evolution of 16 to 32-fold increased vancomycin minimum inhibitory concentration (MIC). To identify the causal genetic variants, we genome sequenced both endpoint resistant clones and whole populations at multiple time-points during the evolution experiment, and reintroduced key mutations, observed in the evolved resistant lineages, into the wild-type ancestral genetic background. Evolution of increased vancomycin resistance was associated with mutations in two distinct pathways, occurring either in *vanT* within the *vanG* cluster or in a gene encoding a regulator of a previously uncharacterised D,D-carboxypeptidase. Mutations in either pathway are predicted to modify the vancomycin target in the cell wall peptidoglycan and both were associated with defects in growth and sporulation of varying severity. Together our results propose a new mechanistic model for vancomycin resistance emergence in *C. difficile*, potentially expanding the genetic determinants of resistance that should be monitored in clinical genomic epidemiology. Moreover, our data suggest that the initial emergence of vancomycin resistance in *C. difficile* in the clinic may be severely constrained by pleiotropic fitness trade-offs with key transmission and virulence traits.

## Results

### Vancomycin resistance evolves rapidly in *C. difficile* during *in vitro* experimental evolution

We first generated genetically barcoded ancestral strains in an avirulent background with either wild-type or elevated mutation rates. Specifically, R20291, a clinically relevant ribotype 027 *C. difficile* strain, was rendered avirulent through complete deletion of 18 kb spanning the entire PaLoc that includes the genes encoding both major toxins and associated regulatory proteins, creating strain R20291Δ*PaLoc*. A subsequent deletion, removing the *mutSL* genes encoding a DNA-damage repair system, generated a hypermutable variant R20291Δ*PaLoc*Δ*mutSL,* with an approximately 20-fold higher mutation rate than the wild-type. Five distinct derivatives of each ancestral strain were then generated through introduction of a 9-nucleotide barcode sequence downstream of the *pyrE* gene, resulting in 10 individually barcoded replicate lines used in the evolution experiment (R20291Δ*PaLoc pyrE*::barcode 1-5; R20291Δ*PaLoc*Δ*mutSL pyrE*::barcode 7-11). Each of the 10 barcoded strains was used to inoculate a 6-well plate containing media supplemented with vancomycin at 0.25x, 0.5x, 1x, 2x, 4x, and 8x the initial MIC of 1 µg/ml. Populations were passaged every 48 h, whereby cells from the well with the highest antibiotic concentration permitting growth were propagated in a 1:400 dilution to a fresh 6-well plate. This process was repeated for a total of 30 serial transfers per replicate line, with adjustment of the vancomycin gradient over time as the growth-permitting vancomycin concentration in each evolving line increased. Ten corresponding control populations were propagated under equivalent conditions in the absence of vancomycin. Populations underwent approximately 8.64 generations per transfer, yielding approximately 259 generations throughout the course of the experiment.

Resistance evolved rapidly in all 10 replicate populations propagated with vancomycin selection (Fig. 1a). Nine of the ten replicate lines evolved to grow in 2 µg/mL vancomycin (an apparent MIC of 4 µg/mL, the EUCAST breakpoint) by the end of the second passage (P2) and all ten grew in the presence of 8 µg/mL (Bc2, 3, 4) or 16 (Bc1, 5, 7, 8, 9, 10, 11) µg/mL vancomycin by P30 (Fig. 1a). This was significantly accelerated in the hyper-mutable replicate lines (Fig. 1b). At least six individual clones were isolated from each evolved population at the end-point and their vancomycin MIC was determined. Of the 82 clones tested, 38 (46%) had an MIC consistent with the vancomycin concentration permitting growth of the population from which they were isolated, while the remainder had an MIC slightly lower than expected [for 42 clones their MIC was 2-fold lower, whereas for 2 clones their MIC was 4-fold lower than the vancomycin concentration permitting growth of the population from which they were isolated (Supplementary Data 1)], demonstrating that significant variation existed within evolved populations by P30.

**Fig. 1.**
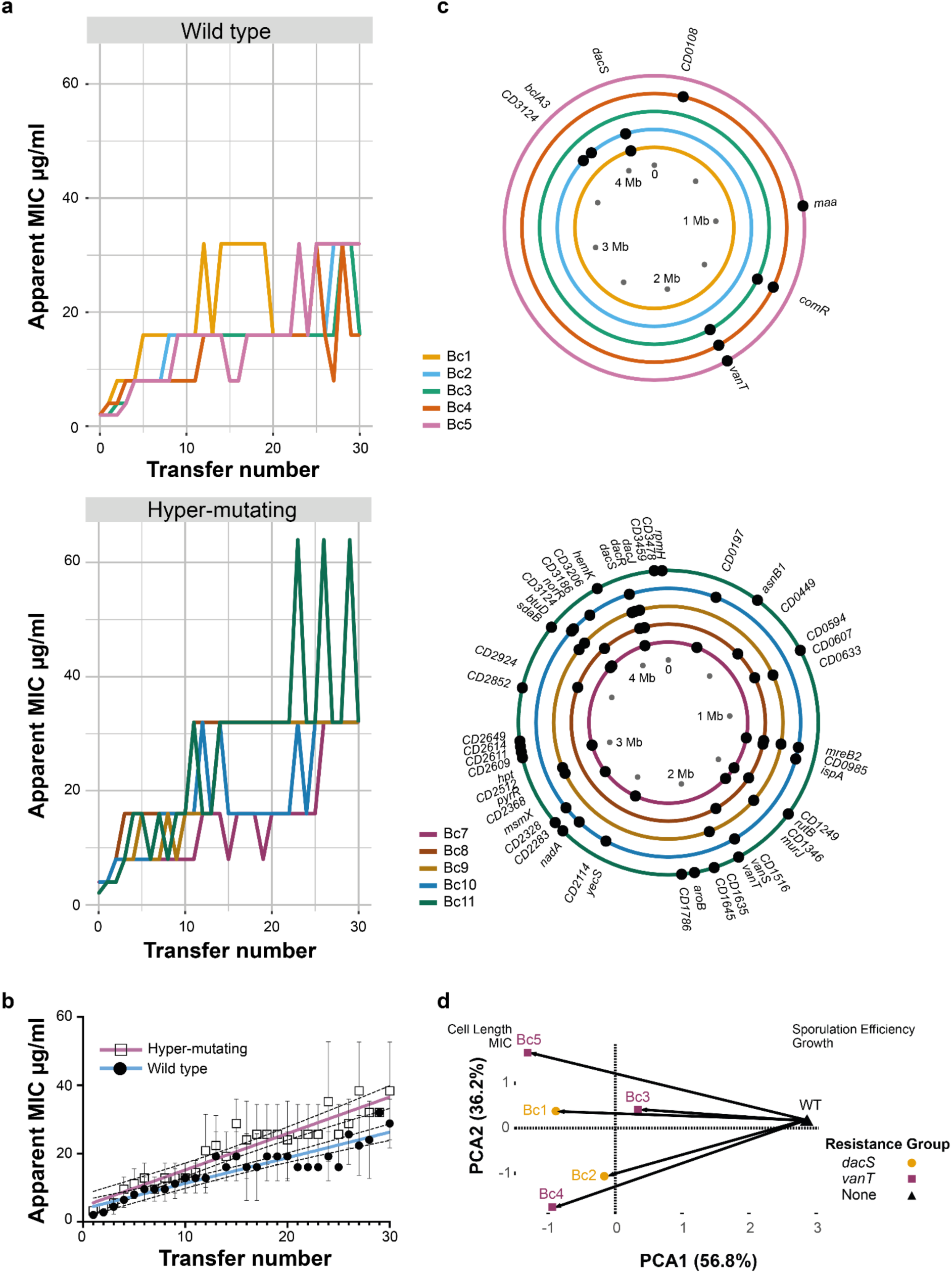
Evolution of vancomycin resistance. **a** Changes in apparent vancomycin MIC over the course of a 30 transfer experimental evolution. MIC was determined as the well with the lowest vancomycin concentration showing no clear growth. **b** Shown are the means and standard deviations of the apparent MIC for five wild type (open squares) and five hyper-mutating (closed circles) populations. Linear regressions fitted to each data set, blue and pink respectively, are significantly different by ANCOVA, P=0.0008. **c** Shown are the chromosomal locations of non-synonymous variant alleles in isolated wild type (top) and hypermutating (bottom) *C. difficile* end-point clones, excluding any mutations that were also identified in any of the control strains. Each circle represents a single *C. difficile* genome, colour coded according to population as indicated in the key on the left. A full list of all variants shown here and including synonymous and intergenic mutations is included in Supplementary Data 2. **d** Principal Component Analysis (PCA) of P30 isolates from populations Bc1-5 (coloured points) vs the ancestral strain (black triangle), with PC1 versus PC2 plotted, accounting for 93% variance. The loadings (sporulation efficiency, growth, MIC, cell length) are shown in respective locations. Arrows show the evolutionary trajectories of wild-type replicate lines from their ancestor in multivariate phenotype space.

### Genetic bases of evolved vancomycin resistance

To understand the genetic bases of increased vancomycin resistance in individual end-point clones, one clone per replicate population that had an MIC representative of their population was selected for whole genome sequencing. The parental strains for each barcoded lineage, and one random endpoint clone from each replicate control population, were also sequenced. For analyses we focused on mutations that were observed in the vancomycin treated lines but never in ancestral or in control evolved clones because these are the most likely to have evolved in response to vancomycin selection. Sequence variants unique to the vancomycin-evolved clones were identified using Varscan^18^, validated with Breseq and IGV^19^ after mapping to the reference R20291 genome (Fig. 1b, Supplementary Data 2). We observed between 1 and 3 unique mutations per genome in wild-type clones and between 14 and 26 in hypermutator clones from the vancomycin treatment. Within gene SNPs accounted for 43% of the variants, of which 67.4% were nonsynonymous and 32.6% were synonymous. Frameshifts accounted for a further 31% of identified unique mutations.

Parallel evolution, where mutations affecting the same locus arise in multiple independently evolving replicate populations, is strong evidence for the action of selection and suggests a potential role for these mutations in adaptation. In the 5 wild-type vancomycin selected lines we observed parallel evolution occurring at three genomic loci: *vanT* in 3 clones, *CD3437* in 2 clones and *comR* in 2 clones. Whereas mutations in *vanT* and *CD3437* were mutually exclusive in wild-type evolved clones, mutations in *comR* always co-occurred with mutations in *vanT.* VanT is a putative Serine racemase (mutations in Bc3-5) encoded within a VanG-type cluster^14^ that was previously implicated in decreased vancomycin susceptibility in *C. difficile*^16^ (Supplementary Data 2). *CD3437* encodes a predicted two-component system histidine kinase (mutations in Bc1 and 2), with its cognate response regulator encoded by *CD3438*. These genes had not previously been implicated in vancomycin resistance, however the nearby *CD3439* encodes a putative D,D-carboxypeptidase that likely plays a role in modification of peptidoglycan through removal of the stem peptide terminal D-Ala. Based on these predicted functions we propose renaming these genes *dacS* (*CD3437*, histidine kinase), *dacR* (*CD3438*, response regulator) and *dacJ* (*CD3439*, D,D-carboxypeptidase). Consistent with mutations at these loci playing key roles in vancomycin resistance, all five hypermutator replicate lines had mutations in the *vanG* operon (1 had a nonsynonymous mutation in *vanT* and 1 in *vanS*) or *dacS*, *dacR* (encoding the cognate response regulator) and *dacJ*. Interestingly, one strain (Bc8) had mutations in both *dacS* and *CD1523* (*vanS*), encoding a two-component system sensor histidine kinase that is thought to regulate the *vanG* operon in response to vancomycin, suggesting that the two pathways to resistance are not entirely mutually exclusive. *comR* encodes a homologue of the RNA degradosome component PNPase, suggesting that RNA stability may play a role in the *vanT*-associated mechanism of vancomycin resistance. Consistent with this possibility, in the third clone carrying a *vanT* mutation we observed coexisting mutations affecting *maa*, encoding a putative maltose O-acetyltransferase, and a 75 bp deletion that completely removed an intergenic region downstream of *rpmH* and before *rnpA*, encoding another predicted component of the RNA degradosome. By contrast, in the 2 clones carrying mutations in *dacS* we did not observe any mutations likely to affect RNA stability: one clone had no additional unique mutations, while the other had nonsynonymous mutations in *bclA3* and *CD3124*. *bclA3* encodes a spore surface protein with no known function in vegetative cells, while *CD3124* encodes an orphan histidine kinase of unknown function. Interestingly, four of the hypermutating lineages also had mutations in *CD3124*, an identical frameshift mutation in all four.

### Vancomycin resistant clones display reduced fitness

To assess the wider consequences of evolved vancomycin resistance for bacterial phenotype, endpoint clones were phenotypically characterised for growth *in vitro*, sporulation efficiency and cell morphology. All 10 strains displayed significantly impaired growth in rich liquid media (Supplementary Fig. 1), with particularly severe defects apparent for Bc10 and 11. Several strains were also impaired in sporulation (Supplementary Fig. 2), with a wide range of phenotypes apparent, from a mild defect for Bc2 and 4 and delayed sporulation for Bc8, to more severe defects for Bc7, Bc9 and 10 and complete loss of sporulation for Bc11. These growth and sporulation defects were also accompanied by changes in cell length relative to the parental wild type strain (Supplementary Fig. 3). Principal component analysis of the five wild type-derived endpoint evolved clones (Fig. 1c) revealed all five resistant strains followed similar evolutionary trajectories away from the parental wild type, associated with lower sporulation efficiency and growth defects, albeit with divergence among replicate lines in the extent of defects and cell size. However, there was no apparent sub-clustering by resistance mechanism.

### Population dynamics in evolving populations

Together the genome sequence data for end-point clones suggests that there are two alternative mechanisms of vancomycin resistance, one involving *dacS* and the other involving *vanT*. To better understand how selection acted upon these mechanisms we next performed pooled population sequencing at passage 10, 20 and 30 to track mutation frequencies over time (Fig. 2 and Supplementary Figs. 4 and 5). In total, discounting variants found in Bc1 P20, we identified 535 unique variants across the 10 parallel populations and three timepoints. We have removed Bc1 P20 from this analysis as the additional 520 variants identified in that sample alone likely reflect random mutation due to the emergence of a spontaneous hyper-mutator phenotype. Impacted genes clustered within 17 distinct functional classes by KEGG analysis (Supplementary Fig. 6), with two component systems and ABC transporters being particularly well-represented. Focusing on the two main routes to resistance identified in endpoint clones, these data revealed highly contrasting selection dynamics, particularly in our wild-type replicate populations: mutations in *dacS* rapidly rose to high frequency, reaching fixation by P10. By contrast, mutations in *vanT* arose later and only reached fixation by P20 or 30, and were preceded by mutations at other sites which reached high frequency by P10 but that ultimately did not survive, being replaced by *vanT* mutants presumably conferring higher levels of vancomycin resistance. The two preceding high frequency mutations in Bc3 (both T>TA) were very close together, separated by only 7 bp in an intergenic region downstream of *CD0482*, encoding a uridine kinase, and approximately 250 bp upstream of *glsA,* encoding a glutaminase. These mutations are outside of the likely promoter region^20^ but it is possible that they affect regulation of *glsA*. Interestingly, changes in glutamine metabolism have previously been linked to vancomycin resistance in *Staphylococcus aureus*^21^. The single high frequency mutation in Bc5 at P10 is a nonsynonymous substitution in *CD3034*, introducing a Gly255Asp mutation in the encoded D-hydantoinase which may play a role in the synthesis of D-amino acids. Taken together, these data suggest that *vanT* is not required for first-step resistance; *vanT* mutations either provide higher-level vancomycin resistance, allowing them to supplant earlier mutations, or they require potentiating mutations to arise and be selected first. However, no consistent secondary mutations common to all populations with *vanT* mutations were identified.

**Fig. 2.**
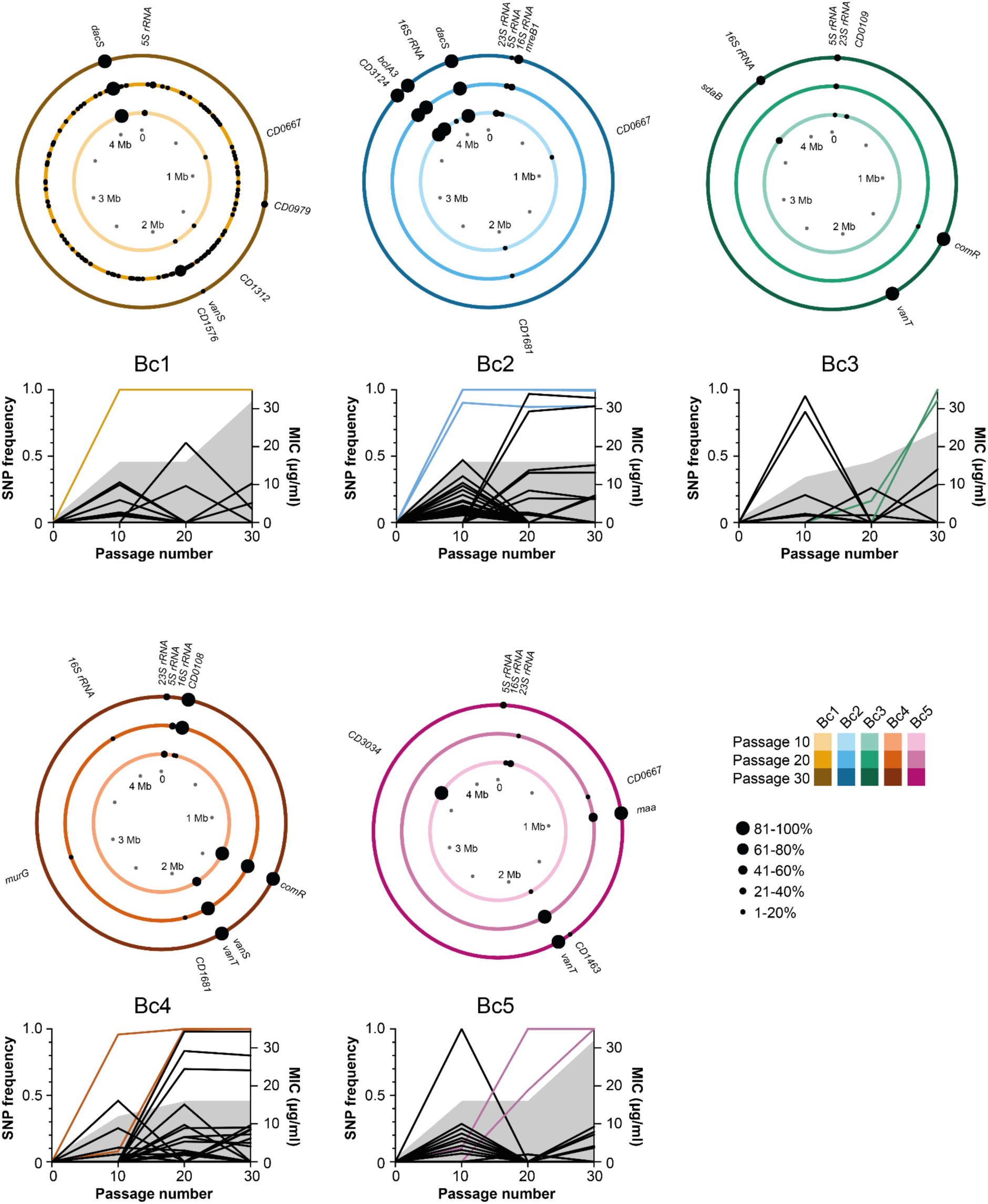
Genomic location of gene variants over time. Accumulation of variants in the wild type *C. difficile* lineages. Each circle plot represents the 4.2 Mb genome of a single evolving population after 10 (inner ring), 20 (middle ring) and 30 passages (outer ring), with the locations of non-synonymous within gene variants indicated with black circles and the penetrance of each mutation in the population indicated by the size of the circle. The line graphs show the frequency of all variants (intergenic, synonymous, non-synonymous, frameshifts and nonsense) in each population. The vancomycin MIC for each population is also indicated by the shaded region. Mutations also identified in the respective end point clone (Fig. 1b) are highlighted by the coloured lines. Note population Bc1 evolved an apparent hypermutator phenotype prior to P20, with 520 variants identified at that time point. For simplicity only variants present in P10 and P30 are labelled. A full list of all variants shown here, including those in Bc1 P20, is included in Supplementary Data 3.

### Recapitulation of *dacS* mutations confirms role in resistance

As the *dacJRS* cluster had not previously been implicated in vancomycin resistance, we validated the role of DacS in vancomycin resistance by recapitulating individual mutations in a clean genetic background. We chose the variant *dacS* alleles identified in Bc1 (714G>T), as this is the sole unique mutation found in that strain, and the variant allele that evolved in parallel in Bc8 and Bc9 (548T>C). Recapitulated strains carrying only the *dacS* mutation of interest were constructed in the parental R20291Δ*PaLoc* by allelic exchange. Introduction of either mutation alone resulted in a 4-fold increase in the vancomycin MIC compared to the parental strain, confirming that DacS is playing a significant role in the evolved resistance we observed (Fig. 3a). AlphaFold prediction of the DacS structure yields a plausible dimer model (Fig. 3b) that is highly similar to previously characterised histidine kinases^22^. The Bc1 and Bc8/9 mutations described here both result in amino acid changes in the DacS cytoplasmic domain: Bc1 Glu238Asp within the predicted catalytic ATPase (CA) domain and Bc8/9 Val183Ala within the dimerization and histidine phosphorylation (DHp) domain. The impact of these mutations on the function of DacS is not clear but we hypothesised that DacS, along with its cognate response regulator DacR, could be regulating the expression of the D,D-carboxypeptidase encoded by *dacJ*. To examine this possibility, we extracted RNA from R20291Δ*PaLoc*, R20291Δ*PaLoc dacS*c.714G>T and R20291Δ*PaLoc dacS*c.548T>C, both in the absence and presence of 0.5 µg/ml vancomycin, and assessed the expression of *dacS*, *dacR* and *dacJ* by qRT-PCR (Fig. 3c). Either point mutation resulted in a dramatic 4.8-94.6-fold upregulation of expression of all three genes and this effect was independent of vancomycin. It is highly likely, therefore, that the overexpression of DacJ and the resulting reduction in vancomycin binding sites within the cell wall accounts for the reduction in vancomycin susceptibility in both strains with mutated *dacS*. The genomic organisation in this region (Fig. 3d) and previous global transcription site mapping^20^ suggests that *dacS* and *dacR* are transcribed from a single promoter upstream of *dacR* and that *dacJ* is transcribed from its own separate promoter. Our data demonstrate that both of these promoters are subject to regulation by the DacS/DacR two component system, although it is not clear if the observed effects are a result of constitutive activation of a TCS that positively regulates these promoters or deactivation of a repressor (Fig. 3d).

**Fig. 3.**
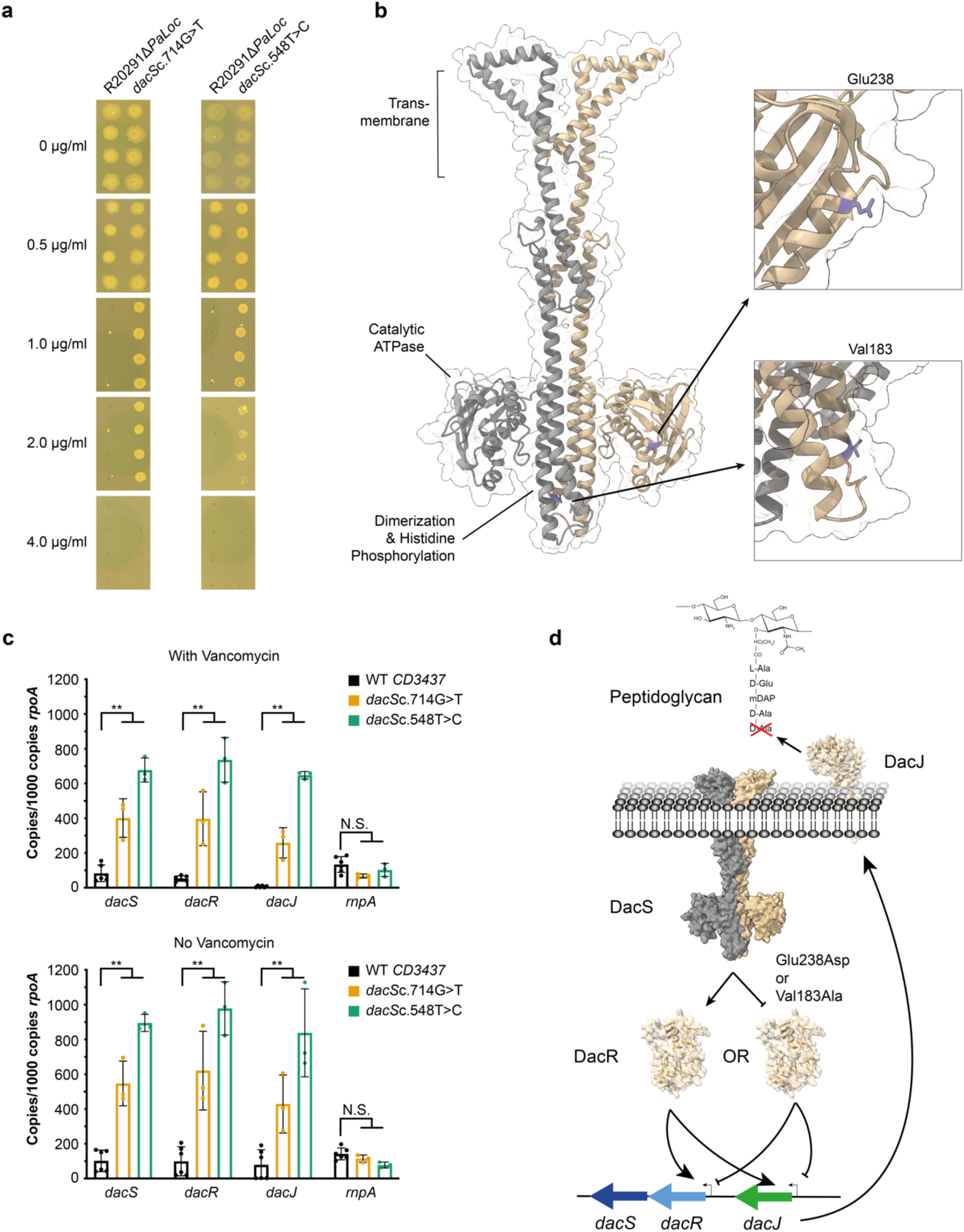
*dacS* mutations lead to dysregulation of *dacJRS*. **a** Vancomycin MICs of R20291Δ*PaLoc*, R20291Δ*PaLoc dacS*c.714G>T and R20291Δ*PaLoc dacS*c.548T>C as determined by agar dilution. Assays were performed in biological triplicate and technical duplicate, four representative spots are shown for each strain. **b** AlphaFold model of DacS as a dimer^23^. The transmembrane domains were identified using DeepTMHMM^24^ and the Catalytic ATPase and Dimerization and Histidine Phosphorylation domains were predicted using InterProScan^25^. The locations of Val183 and Glu238 are highlighted in purple on one chain. **c** qRT-PCR analysis of *dacJRS* expression in R20291Δ*PaLoc* (black bars), R20291Δ*PaLoc dacS*c.714G>T (yellow bars) and R20291Δ*PaLoc dacS*c.548T>C (green bars). *rnpA,* which is implicated in the *vanT* resistance pathway, was included as an additional unrelated control. Expression was quantified against a standard curve and normalised relative to the house-keeping gene *rpoA*. Assays were performed in biological and technical triplicate. Statistical significance was calculated using a two-way ANOVA with Dunnett’s multiple comparison, ** = P<0.001. **d** Two mechanisms by which DacS mutations could alter expression of *dacJRS*. Phosphorylated-DacR could be an activator or repressor of the two promoters in the *dacJRS* locus, with DacS Glu238Asp or Val183Ala substitutions either constitutively activating or inhibiting the function of the TCS respectively. In either scenario, the consequence is over-expression of DacJ which is then translocated to the cell surface where it can cleave the terminal D-Ala residue in nascent peptidoglycan, thereby preventing vancomycin binding.

## Discussion

Vancomycin is one of the few antibiotics in routine use for treatment of CDI worldwide and is now the frontline drug of choice in the UK^7^. High level vancomycin resistance is widespread in *Enterococcus* spp. and in *S. aureus* but has yet to be reported in *C. difficile*, where there are few verifiable reports of reduced susceptibility in clinical strains, despite anecdotal reports of vancomycin treatment failure^26^. However, it is not clear if this apparent lack of resistance reflects an underlying constraint upon the emergence of resistance in this species or is simply an artefact of a lack of routine monitoring in the clinic. Here we show using laboratory experimental evolution that *C. difficile* can rapidly evolve high-level vancomycin resistance via two alternative mechanisms, but that increased resistance is associated with severe pleiotropic effects, including growth and sporulation defects, which may act to limit the emergence of resistance in clinical settings.

Under ramping vancomycin selection, resistance emerged rapidly in all 10 replicate lines reaching 16 to 32-fold higher MIC within approximately 250 generations. Whole genome sequencing of individual clones from each population at the end of the evolution revealed two evolutionary pathways to resistance, centring around mutations in *vanT*, encoding the Serine/Alanine racemase component of a VanG-type cluster, and mutations in a cluster of genes encoding a two component system and a D,D-carboxypeptidase (*dacJRS*). The VanG cluster is common in *C. difficile* strains but its potential role in vancomycin resistance had been disputed^14,15^. More recently however, mutations in the VanRS two-component system that derepress the rest of the cluster and reduce vancomycin susceptibility have been identified in both laboratory evolution experiments and in clinical isolates^16,17^. These observations confirm that changes to the expression of genes in the VanG cluster can indeed contribute to reduced vancomycin susceptibility. Interestingly we detected *vanS* mutations only transiently in our evolution and none became fixed, suggesting they had only a limited contribution to resistance or were accompanied by severe fitness defects (Fig. 4). In contrast, mutations to *vanT* were common, fixing in four of ten replicate lines. Mutations in the *dacJRS* cluster were also extremely commonly observed and often rose to high frequency, with variant *dacS* alleles fixing in three populations (Bc1, 2 and 9), an identical *dacR*c.532A>G variant fixing in two populations (Bc7 and 10) and a *dacJ* variant fixing in Bc9. Mutations in *dacS* and *dacR* also transiently fixed in two further populations (Bc8 and 10 respectively) at P20 before being subsequently outcompeted. Interestingly the *dacS* mutation identified in Bc8 at P20 (548T>C) is identical to that observed in endpoint isolates from Bc9. None of these genes had been previously linked to vancomycin resistance but the predicted D,D-carboxypeptidase activity of DacJ points to a plausible mechanism through removal of the terminal D-Ala residue in nascent peptidoglycan^27,28^ (Fig. 3d). We hypothesised that the two-component system encoded by *dacS* and *dacR* was regulating expression of *dacJ*, providing a mechanistic link for all of these mutations. This was confirmed by recapitulation of two distinct *dacS* mutations (from Bc1 and Bc8/9) in a clean genetic background, leading to derepression of both *dacJ* and the *dacSR* bicistronic operon. Importantly this effect was independent of vancomycin, leaving open the possibility that the wild type system could still contribute to vancomycin resistance in the appropriate permissive environmental condition.

**Fig. 4.**
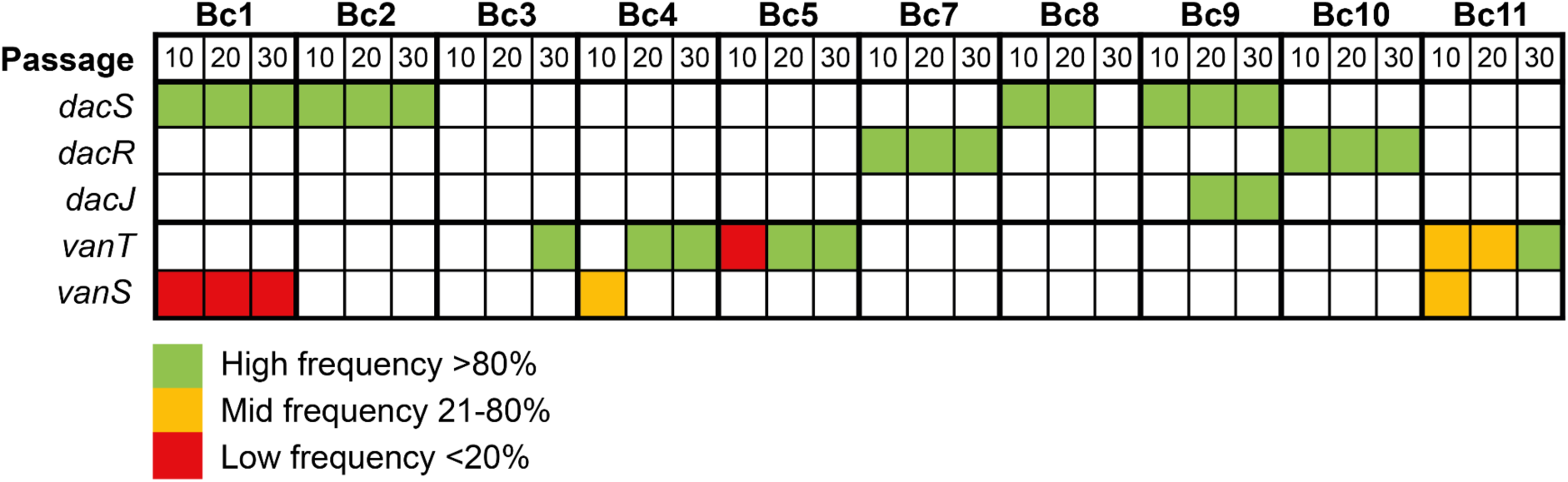
Two distinct pathways to resistance. Relative frequencies and time of emergence of mutations in genes *dacS*, *dacR*, *dacJ*, *vanT* and *vanS* across all ten evolving populations.

Analysis of evolutionary dynamics over the course of the evolution also revealed intriguing differences in the timing of *dacJRS* vs *vanT* mutations (summarised in Fig. 4). *dacJRS* mutations had fixed by P10 in nearly every population in which they persisted to the end of the evolution, the exception being Bc9 *dacJ* which first appeared and fixed at P20. However, that population also had a superseding *dacS* mutation that had fixed by P10. By contrast, mutations in *vanT* typically didn’t fix until P20 or P30, although sometimes present at lower frequency at earlier timepoints. Delayed emergence of *vanT* variants may also indicate a reliance on preexisting potentiating mutations, although no consistent secondary mutations were found in all *vanT* lineages. The two resistance mechanisms also seemed to be mutually exclusive, *vanT* mutations did not co-occur with *dacJRS* in any population or endpoint isolate. It is possible that early emergence of mutations in *dacJRS* precludes subsequent mutations to *vanT* and commits the population to that pathway.

In all populations, emergence of resistance in our experiment was accompanied by severe, if somewhat variable, fitness costs, importantly including severe sporulation defects in two lineages. As the spore is the infectious form of *C. difficile* and an absolute requirement for patient to patient transmission^29^, these sporulation defects would likely have serious consequences for the infectivity and onward transmission of evolved resistant isolates. The genetic basis of the sporulation defects in these isolates is not clear, as sporulation is an extremely complex and poorly understood cell differentiation process^30,31^, but if a similar trait were to emerge in a patient during vancomycin treatment this would be an evolutionary dead end. These fitness defects may well explain delays to the emergence of vancomycin resistance in the clinic. However, it is also possible that the accumulation of refining mutations would eventually lead to gradually improving fitness as has been seen in long term evolution experiments^32,33^. Indeed, we saw extensive evidence of succession here, with early high frequency mutations that conferred moderate increases in MIC being completely supplanted by later variants, of presumably improved fitness. The potential for further evolution in our experiment was also clear from the continual emergence of new variants, even at the final timepoint. It remains to be seen how far towards full resistance *C. difficile* can go given the enough time and the right conditions but, given the increasing reliance on vancomycin in the treatment of CDI, it is crucial that we begin to understand the possible routes to resistance. The question of how likely *C. difficile* vancomycin resistance is in the real world remains open but this work, and other efforts towards understanding possible mechanisms of resistance, will hopefully provide a roadmap to guide genomic surveillance efforts.

## Methods

### Strains and Growth Conditions

All strains used or generated in the course of this study are described in Supplementary Table 1. *C. difficile* was routinely cultured on brain heart infusion (BHI) agar and in tryptone yeast (TY) broth. *C. difficile* was grown in an anaerobic cabinet (Don Whitley Scientific) at 37°C, with an atmosphere composed of 80% N_2_, 10% CO_2_ and 10% H_2_. Media was supplemented with thiamphenicol (15 µg/mL) (Sigma) and colistin (50 µg/mL) (Sigma) as appropriate. For counter selection against plasmids bearing the *codA* gene, *C. difficile* differential media with 5-fluorocytosine (CDDM 5-FC) was used as described previously^34^. *E. coli* was routinely cultured in Luria-Bertani (LB) broth or agar at 37°C, and supplemented with chloramphenicol (15 µg/mL) (Acros Organics) or kanamycin (50 µg/mL) (Sigma) as appropriate.

### Molecular biology

All oligonucleotides and plasmids are described in Supplementary Tables 2 and 3 respectively. Plasmid miniprep, PCR purification and gel extractions were performed using GeneJET kits (Thermo Fisher). High-fidelity PCR amplification was performed using Phusion polymerase (NEB), and standard PCR amplification was performed using Taq mix red (PCRBIO), according to manufacturers’ instructions. Gibson assembly primers were designed using NEBBuilder (NEB) and restriction digestion and DNA ligation was performed using enzymes supplied by NEB. Plasmids were transformed into NEB5a (NEB) or CA434 competent *E. coli* cells according to the NEB High Efficiency Transformation Protocol. The sequences of cloned fragments were confirmed using Sanger sequencing (Genewiz, Azenta Life Sciences, Germany).

### C. difficile mutagenesis

*C. difficile* strain R20291 was modified for use in evolution experiments and to recapitulate individual mutations. Homology arms for introducing mutations onto the *C. difficile* chromosome by allelic exchange were generated either by Gibson assembly of PCR products or synthesised by Genewiz (Azenta Life Sciences) and then subsequently cloned between BamHI and SacI sites in pJAK112^20^, a derivative of pMTL-SC7215^34^. Following confirmation by Sanger sequencing, plasmids were transformed into *E. coli* CA434 and transferred to *C. difficile* by conjugation^35^. Homologous recombination was performed as previously described^34^ and mutations were confirmed by PCR and Sanger sequencing of mutated regions.

R20291 was rendered avirulent via deletion of the complete pathogenicity locus (encoding toxins A and B) using plasmid pJAK143^36^, yielding strain R20291Δ*PaLoc*. This strain was then further modified by deletion of DNA repair operon *mutSL* to create a hyper-mutator strain. 1.2 kb up and downstream of the *mutSL* operon was amplified by PCR using oligonucleotides RF2066 and RF2067, and RF2068 and RF2069 (Supplementary Table 2), respectively, and cloned between BamHI and SacI sites in plasmid pJAK112. The resulting plasmid, pJEB002, was then conjugated into R20291Δ*PaLoc* and allelic exchange performed as described above, yielding R20291Δ*PaLoc*Δ*mutSL*. Enumeration of CFUs on BHI agar containing rifampicin (0.015 µg/mL) suggested this strain has a mutation rate approximately 20-fold higher than R20291Δ*PaLoc*. Five barcoded variants of both R20291Δ*PaLoc* and R20291Δ*PaLoc*Δ*mutSL* were then generated by insertion of unique sequencing barcodes. Briefly, pJAK081 (identical to pJAK080^20^ but in the pMTL-SC7215 backbone) was modified by introduction of a synthetic DNA fragment containing a multiple cloning site and a 9 bp sequencing barcode (Barcode 1), flanked by the *fdx* and *slpA* terminators, between the existing homology arms for insertion of DNA between *CD0188* (*pyrE*) and *CD0189* in the R20291 genome. A second plasmid (pJAK202) containing Barcode 2 was constructed in the same manner and the rest (pJAK203-205 and 207-211) were generated via site directed mutagenesis using pJAK201 as a template. These ten plasmids were then used to generate R20291Δ*PaLoc* Bc1-5 and R20291Δ*PaLoc*Δ*mutSL* Bc7-11. Recapitulated strains were generated by introducing point mutations into R20291Δ*PaLoc.* An approximately 2 kb synthetic DNA fragment, centred on the mutation of interest, was cloned between BamHI and SacI sites in pJAK217, generating pJEB019 (*dacS*c.548T>C) and pJEB026 (*dacS*c.714G>T). The resulting plasmids were conjugated into R20291Δ*PaLoc,* and allelic exchange was performed as described above, generating R20291Δ*PaLoc dacS*c.548T>C and R20291Δ*PaLoc dacS*c.714G>T respectively.

### Evolution

Directed evolution of *C. difficile* was performed using a broth-based gradient approach in which 10 individually barcoded parallel lines were evolved for a period of 30 passages. A 6-well plate for each of the parallel lines was prepared for each passage using 4 mL of TY broth with vancomycin, spanning a gradient of 0.25 to 8x the current MIC, as determined from the most recent passage for each line, allowing the gradient to rise with increasing resistance.

The evolution was initiated using overnight cultures from single colonies, adjusted to OD_600nm_ 0.1. 10 µL was added to each well, before incubating for 48 h at 37°C. Plates were visually inspected after 48 h and 10 µL of the well with the highest antibiotic concentration supporting growth was used to inoculate the wells of the subsequent passage. For each parallel line, a control well was passaged without antibiotic. 1 mL of a population, and the corresponding control, was frozen at -80°C in 15% glycerol whenever the MIC increased; and after passages 10, 20 and 30.

### gDNA Extraction

gDNA was obtained from *C. difficile* cultures using the phenol-chloroform method as described previously^30^. DNA concentration was quantified using Qubit, and purity was assessed via microvolume spectrometry.

### Sequencing

End-point (P30) isolates and respective controls were gained through plating P30 populations and culturing single colonies. Vancomycin MICs were determined by agar dilution (described below), and gDNA extraction and quantification were performed as above. Library prep (Nextera XT Library Prep Kit (Illumina, San Diego, USA)) and 30x illumina sequencing (NovaSeq 6000, 250 bp paired end protocol) of isolates was performed at MicrobesNG (Birmingham, UK). Reads were trimmed at MicrobesNG using Trimmomatic (v0.30)^37^ with a sliding window quality cut-off of Q15.

Pooled sequencing of the 10 evolved and 10 control populations was performed at 3 timepoints – P10, P20 and P30. 1 mL of the well with highest antibiotic concentration supporting growth was taken after 48 h, harvested via centrifugation, and frozen at -20°C. gDNA extraction and quantification were performed as above. Library prep (Nextera DNA Flex Library Prep Kit (Illumina, San Diego, USA)) and 250x Illumina sequencing (NovaSeq 6000, 150bp paired end protocol) was performed at SNPsaurus (Oregon, USA). Reads were trimmed using Trimmomatic (v0.39) with the following criteria: leading:3; trailing:3; slidingwindow:4:15; minlen:36.

### Sequencing Analysis Pipeline: Isolates

Trimmed reads were checked using FastQC (v0.11.9)^38^ to ensure sufficient quality for analysis. Analysis was primarily performed using a custom script, built based on a resistant mutant analysis pipeline^39^. Reads were aligned to the reference (*C. difficile* R20291, accession number: FN545816) using BWA-mem (v0.7.17)^40^ and sorted using SAMtools (v1.43)^41^. Coverage across the genome was inferred using Bedtools (v2.30.0)^42^ genomecov and map functions. PCR duplicates were removed via Picard (v2.25.2) (http://broadinstitute.github.io/picard/). The mpileup utility within SAMtools (v1.43) was used to generate the mpileup file required for Varscan. Variants were then called using Varscan (v2.4.3-1)^18^ mpileup2cns (calling SNPs and indels) using the following parameters: min-coverage 4; min-reads2 4; min-var-freq 0.80; p-value 0.05; variants 1; output-vcf 1. Here, a minimum of 4 reads were required to support a variant, and the cut-off for variant calling was 80%. Vcf files were annotated using snpEff (v5.0)^43^. Variants were also called using Breseq (v0.35.5)^19^, using default parameters, and putative variants were retained if detected in both analysis pipelines. All variants were manually verified using IGV (v2.8.6)^44^. Variants that were also called in control lines were removed using Varscan (v2.4.3-1) compare, generating a list of variants unique to vancomycin resistant lines.

Nonsynonymous mutations occurring within genes were plotted using a previously published custom script in RStudio (v4.1.0) utilising the Plotrix package^45^ to visualise parallel evolution.

### Sequencing Analysis Pipeline: Populations

The 10 evolved and 10 control populations, sequenced at P10, P20 and P30, were evaluated using a population analysis pipeline that followed the same custom script as for isolate analysis, with modifications to SNP calling parameters and filtering. InSilicoSeq (v1.5.4)^46^ was used to rapidly simulate realistic sequencing data at a range of coverage depths (80x, 100x, 300x) based on the R20291 genome, with SNPs seeded at 5% frequency. Simulated sequences were analysed using the custom pipeline, and a range of Varscan parameters (P-values, min-coverage, min-reads2) were trialled to generate a set of parameters to accurately call variants without false positives.

Varscan parameters used to call population variants were dependent on average coverage depth, allowing a strong evidence base whilst calling low frequency variants. Samples were separated by average coverage: the following Varscan (v2.4.3-1) mpileup2cns parameters were used for samples with 100x or higher average coverage: min-coverage 80; min-var-freq 0.05; p-value 0.05; variants 1; output-vcf 1. A minimum of 4 reads were required to support a variant, and the cut-off for variant calling was 5%. For samples with average coverage below 100x, the following Varscan (v2.4.3-1) mpileup2cns parameters were used: min-coverage 4; min-reads2 4; min-var-freq 0.05; p-value 0.05; variants 1; output-vcf 1. As with high coverage samples, a minimum of 4 reads were required to support a variant, meaning the minimum variant frequency which could be called was inversely scaled (4/coverage at position), with a minimum frequency of 5%. This process identified a list of variants within each line at each time point, and their respective frequencies.

As in the isolate pipeline, population variants were filtered using Varscan (v2.4.3-1) compare, removing variants that were also present in control lines to generate a list of variants unique to vancomycin resistant lines. Further filtering of this dataset, again using Varscan (v2.4.3-1) compare, discarded variants that never reached above 10% frequency by the end of the evolution. Variants were also manually inspected to ensure no variants remained the same frequency across all time points.

Mutations occurring within genes (except those in Bc1 P20) were coloured according to their barcoded replicate line, and displayed in KEGG Mapper - Color^47^ to view KEGG pathways. Nonsynonymous mutations occurring within genes and rRNAs were plotted using a previously published custom script in RStudio (v4.1.0) utilising the Plotrix package^45^ to visualise population dynamics, where point size indicated mutation frequency in the population.

### Strain fitness analysis

Growth curves were performed anaerobically in 96 well plates using a Stratus microplate reader (Cerillo). Overnight cultures of *C. difficile* were diluted to OD_600nm_ 0.05 and incubated at 37°C for 1 h. Plate lids were treated with 0.05% Triton X-100 + 20% ethanol. Plates were prepared using 200 µL of equilibrated culture. Samples were measured at minimum in biological and technical triplicate. The OD_600nm_ was measured every 3 min over a 24 h period. Data was plotted in Graph pad Prism (v9.0.2), and was analysed in RStudio (v4.1.0) using the GrowthCurver package (v0.3.0)^48^.

To assess sporulation efficiency, triplicate overnight *C. difficile* cultures were adjusted to OD_600nm_ 0.01 and grown for 8 h. Cultures were then adjusted again to OD_600nm_ 0.01, subcultured 1:100 into 10 mL BHIS broth, and grown overnight to obtain early stationary phase spore-free cultures (T = 0). At T = 0, and the following 5 days, total viable counts were enumerated by spotting 10-fold dilutions in technical triplicate onto BHIS agar with 0.1% sodium taurocholate. Colonies were counted after 24 h incubation. Spore counts were enumerated using the above method following heat treatment (65°C for 30 min).

To visualise cell morphology, *C. difficile* samples were harvested via centrifugation, washed twice in PBS, and fixed in 4% paraformaldehyde; before harvesting and resuspension in dH_2_O. Samples were mounted in 80% glycerol, and imaged using a 100x Phase Contrast objective on a Nikon Ti eclipse widefield imaging microscope using NIS elements software. Images were analysed in Fiji (v2.9.0) using MicrobeJ (v5.13l)^49^.

### Vancomycin MICs

MICs were obtained via standard agar dilution methods^50^. Briefly, overnight cultures were adjusted to OD_600nm_ 0.1. 2.5 µL of sample was spotted in biological triplicate and technical duplicate onto BHI plates with ranging antibiotic concentrations. MICs were determined after 48 h incubation, and plates were imaged using a Scan 4000 colony counter (Interscience).

### Principal Component Analysis

WT (Bc1-5) P30 resistant isolates were characterised in terms of their sporulation efficiency, growth rate, MIC and cell length. These were compared, along with the WT ancestor from the start of the evolution, in a principal component analysis (PCA). The PCA was computed using the prcomp() function in Base R (http://www.rproject.org/), and visualised using the factoextra package. The first 2 PC were plotted, as these accounted for 93% variance. The loadings were added in their respective locations, and isolates were coloured based on resistance mechanism.

### qRT-PCR

Total RNA was extracted using the FastRNA pro blue kit (MP Biomedicals). Cells were grown to an OD_600nm_ of approx. 0.4, and 2 volumes of RNA protect (Qiagen) were added. Cells were incubated for a further 5 min, before harvesting via centrifugation. Cell pellets were stored at -80°C. Pellets were resuspended in RNA Pro solution, and transferred to a tube containing lysing matrix B (MP Biomedicals). Cells were lysed via FastPrep (2 cycles of 20 s, 6 m/s), and centrifuged (16,200 x g, 4°C, 10 min) to remove insoluble cell debris. The supernatant was transferred to a microfuge tube, and 300 µL chloroform (Sigma) was added. Samples were centrifuged again (13,000 rpm, 4°C, 15 min), and the supernatant was precipitated at 20°C overnight after addition of 500 µL 100% ethanol. After precipitation, RNA was harvested by centrifugation (13,000 rpm, 4°C, 15 min), washed with 70% ethanol, and dried. Precipitated RNA was resuspended in 50 µL nuclease-free water, residual DNA was removed using the Turbo DNA-free kit (Invitrogen) and the RNA was cleaned and concentrated with the RNeasy Minelute cleanup kit (Qiagen), as per manufacturers’ instructions.

cDNA was generated using Superscript III (Invitrogen). 5 µg RNA was mixed with 2 µL dNTP mix and 1 µL 100 mM random primer (Eurofins), heated at 65°C for 5 min, and cooled on ice. 8 µL 5X buffer, 2 µL 0.1M DTT (Invitrogen), 1 µL RiboLock RNase inhibitor (Thermo Scientific) and 2 µL Superscript III were added, before incubation at 25°C (5 min), 50°C (30 min) and 70°C (15 min). cDNA was adjusted to 40 ng/µL. RT negative controls were made as above, without presence of Superscript III.

Expression was measured against an exact copy number control, via standardisation with a plasmid containing one copy of each target DNA sequence^51^. *rpoA* was used as the housekeeping gene to normalise results, and *rnpA* was used as an additional unrelated control. A plasmid (pJEB029) was synthesised with the pUC-GW-Kan backbone (Genewiz), containing approximately 200 bp target gene fragments of *dacS, dacR, dacJ*, *rpoA* and *rnpA.* This was purified using the GeneJET plasmid miniprep kit (Thermo Fisher), and linearised using NdeI (NEB) and diluted to known copy number (2 × 10⁸ per µL). A qPCR mastermix was assembled, containing 25 µL SYBR Green JumpStart Taq ReadyMix (Sigma), 7 µL MgCl2 in buffer (Sigma), forward and reverse primers (concentration determined by prior optimisation) and nuclease-free water up to 45 µL. 5 µL of cDNA (40 ng/µL), 5 µL of RT negative control, or 5 µL qPCR plasmid pJEB029 (diluted serially in lambda DNA (Promega)) were added, and qPCR was performed (BioRad CFX Connect Real Time System). Copy numbers were calculated using BioRad CFX manager (v3.1), and data were analysed in Microsoft Excel (2016) to generate copies per 1000 copies of *rpoA*. Data were graphed and statistically analysed in Graph pad Prism (v9.0.2).

### Structural modelling of DacS

DacS was modelled as a dimer using AlphaFold^23^ and the resulting output files were visualised using ChimeraX^52^.

### Statistics

Statistical analysis was performed in Graphpad Prism (v9.0.2), and P <0.05 was considered significant. Data are presented as mean ± SD, unless otherwise stated. Differences in growth were analysed using the R package GrowthCurver outputs. A cross-correlation was performed in RStudio using Hmisc and corrplot packages^53,54^, which determined area under curve (AUC-E) to be the most representative measure of growth. The differences in AUC-E, compared to the control strain curves, were calculated using t-tests with Welch’s correction. To test differences in sporulation efficiency, AUC was chosen as a representative measure of sporulation across all timepoints. AUC was calculated in Graphpad Prism (v9.0.2), and the difference between AUC for P30 isolates was compared with the WT using a one-way ANOVA with Brown-Forsythe and Welch’s correction. Statistical analysis of cell length was performed in Graphpad Prism (v9.0.2). Fiji (v2.9.0) MicrobeJ (v5.13l) cell length outputs for P30 isolates were compared to the WT using a 1-way ANOVA with Brown-Forsythe and Welch’s correction to test for differences in cell length. To determine whether differences in expression of *dacS*, *dacR, dacJ* and *rnpA* were significant between R20291Δ*PaLoc,* R20291Δ*PaLoc dacS*c.548T>C and R20291Δ*PaLoc dacS*c.714G>T, a two-way ANOVA with Dunnett’s multiple comparisons was performed in Graphpad Prism (v9.0.2).

## Data Availability

Genome sequence data for parental strains, P30 resistant isolates and respective controls, as well as pooled population sequencing data for resistant populations and controls at P10, 20 and 30 is deposited with the European Nucleotide Archive (ENA). The accession numbers for these may be viewed in Supplementary Table 4.

## Acknowledgements

We thank the Wolfson Light Microscopy Facility at the University of Sheffield, for assistance with phase-contrast microscopy and the Bioinformatics Core, for training. We would also like to thank Joseph A Kirk for construction of barcoding and Δ*PaLoc* plasmids, and for the R20291Δ*PaLoc* strain. JEB is supported by a studentship from the MRC Discovery Medicine North (DiMeN) Doctoral Training Partnership (MR/R015902/1). RCTW is supported by BBSRC grant (BB/T014342/1) and MAB by a Wellcome Trust Collaborative Award (220243/Z/20/Z). CET is a Royal Society & Wellcome Trust Sir Henry Dale Fellow (208765/Z/17/Z).

## Author contributions

JEB carried out all experiments, collected and analysed data, revised the sequencing analysis pipelines, wrote and revised the manuscript. RCTW provided guidance on sequencing analysis. CET designed experiments, provided guidance and provided tools for qRT-PCR experiments and analysis. RRC provided guidance on bioinformatics analysis, provided bioinformatics scripts, and analysed KEGG data. MAB designed the study, interpreted evolution data, supervised the study, and revised the manuscript. RPF designed the study, analysed data, supervised the study, wrote and revised the manuscript.

## Materials and correspondence

Requests should be addressed to Robert Fagan or Michael Brockhurst.

