## Supplemental Figures and Tables for "Multiple evolutionary pathways lead to vancomycin resistance in *Clostridioides difficile*"

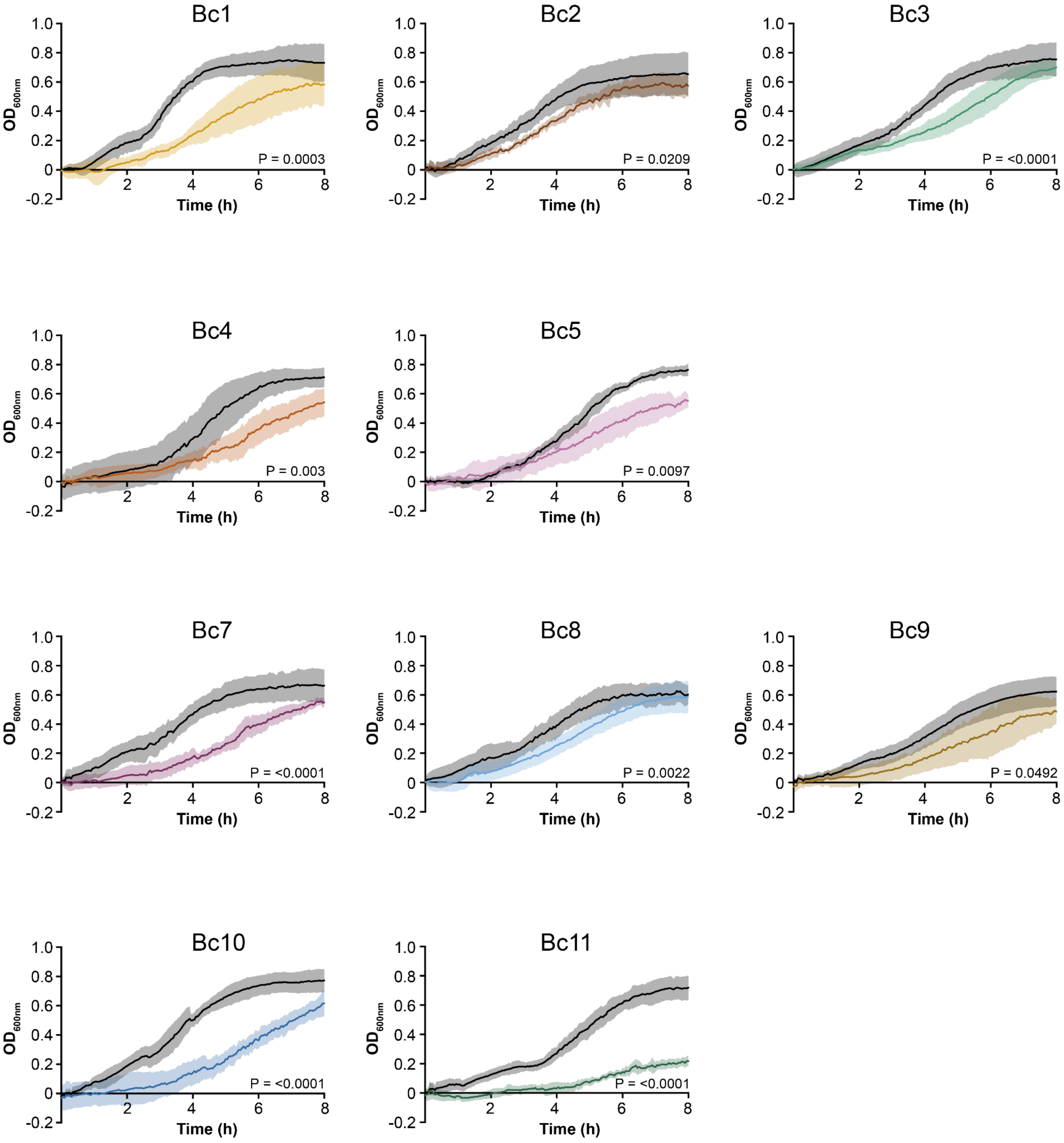


**Supplementary Figure 1. Growth of evolved endpoint clones.** Growth over time in rich media (TY broth) was measured at 600 nm in a 96 well microplate spectrometer. Growth of each endpoint clone (coloured lines) was compared to its matched control (black lines). Shown are the mean and standard deviation of repeats, assayed at minimum in biological and technical triplicate. For each strain, area under the curve was determined using the GrowthCurver R package and these were compared using Student's t-tests with Welch's correction, with the P value shown on each graph. All pairwise differences are significant.

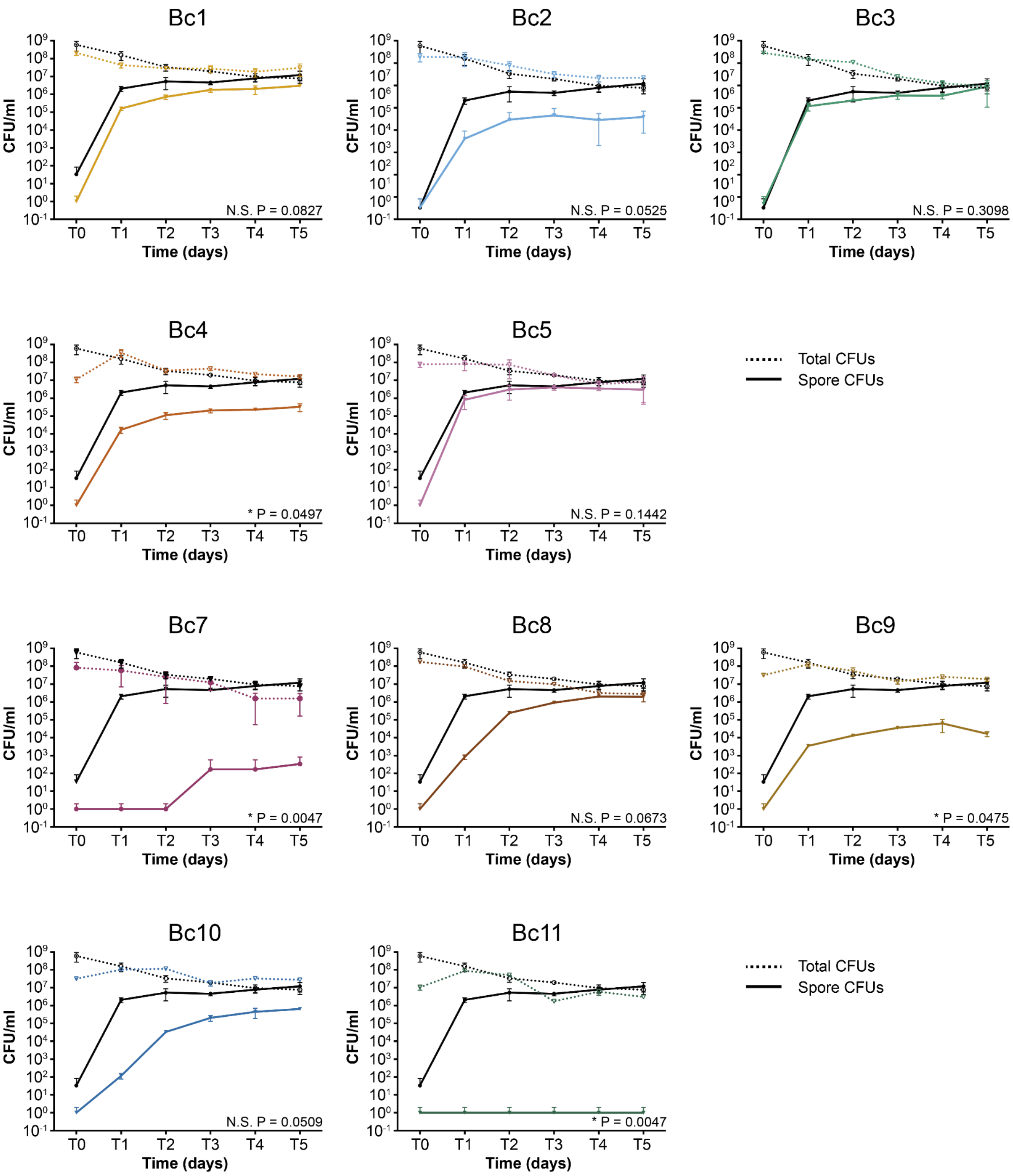


**Supplementary Figure 2. Sporulation of evolved endpoint clones.** Sporulation efficiencies of each endpoint clone (coloured lines) were compared to the parental R20291Δ*PaLoc* (black lines). Stationary phase cultures were incubated anaerobically for 5 days with samples taken daily to enumerate total colony forming units (CFUs, dotted lines) and spores (solid lines), following incubation at 65°C for 30 min to kill vegetative cells. Shown are the mean and standard deviations of biological duplicates assayed in triplicate. For each strain, spore CFU area under the curve was determined using Graphpad Prism and these were compared using Dunnett's T3 multiple comparisons test with the adjusted P value shown on each graph. * = significant difference, N.S. = not significant.

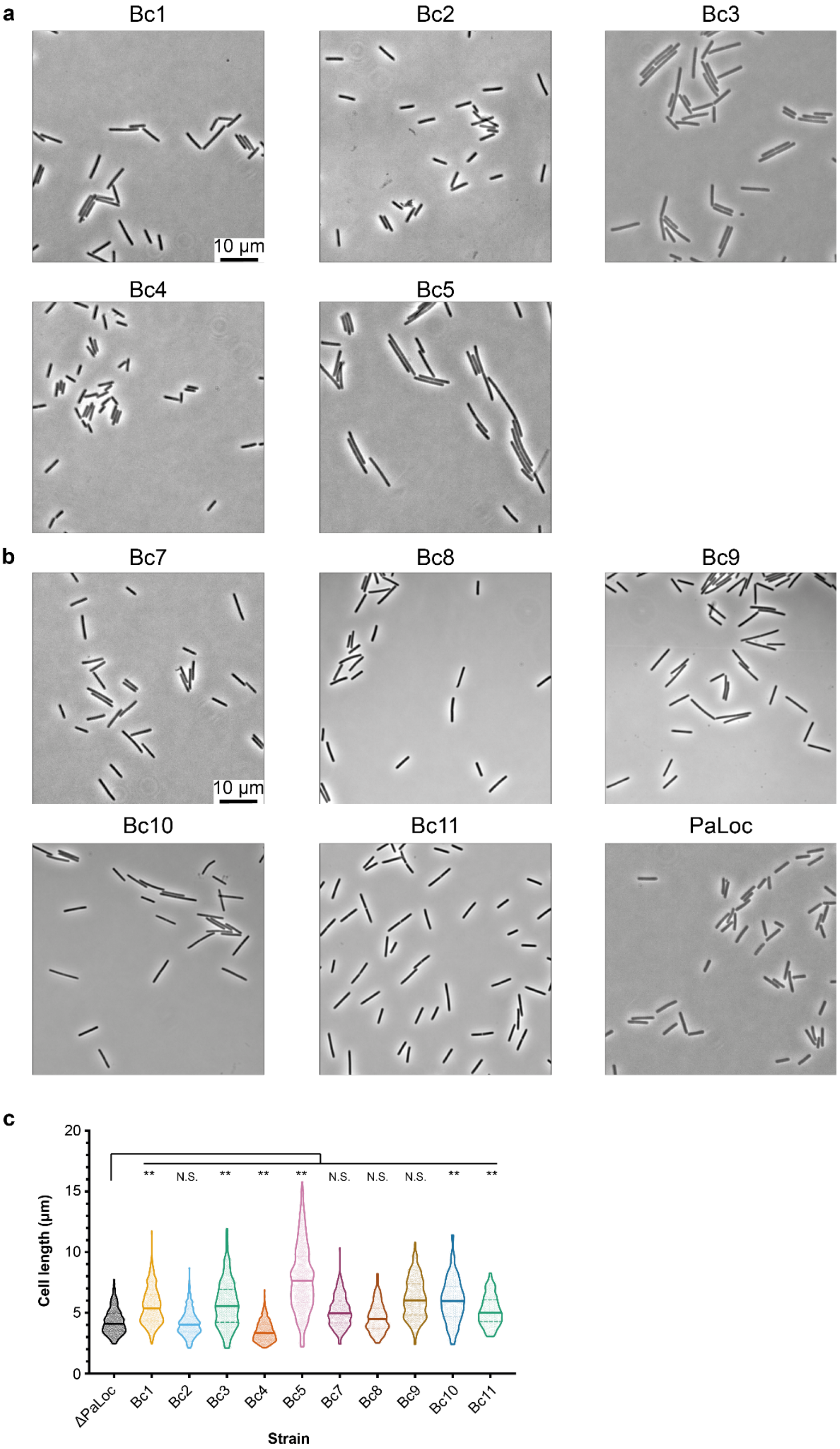


**Supplementary Figure 3. Cell morphology of evolved endpoint clones.** Phase contrast light microscopy of mid-log cultures of each wild type (**a**) and hypermutating (**b**) endpoint clone, with R20291Δ*PaLoc* for comparison. Shown is a representative field of view for each strain. **c**

Images were analysed using MicrobeJ to determine lengths of at least 185 individual cells for each strain. Shown is an all point violin plot with the median indicated by a solid horizontal line. Statistical significance of evolved isolates against the R20291Δ*PaLoc* control was calculated using a one-way ANOVA with Dunnett's T3 multiple comparisons test, ** = P<0.0001, N.S. = not significant.

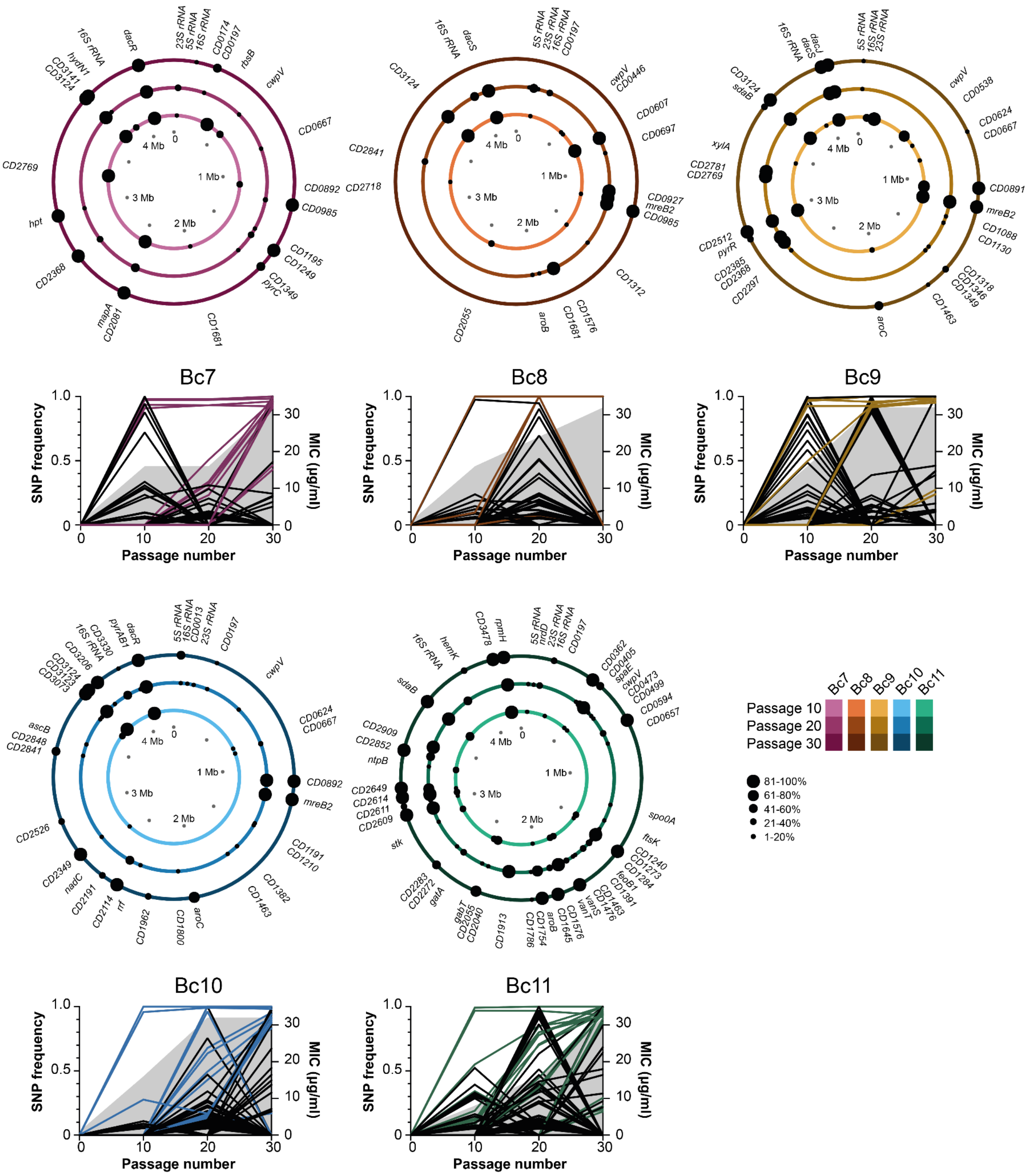


**Supplementary Figure 4. Genomic location of gene variants over time.** Accumulation of variants in the hyper-mutating *C. difficile* lineages. Each circle plot represents the 4.2 Mb genome of a single evolving population after 10 (inner ring), 20 (middle ring) and 30 passages (outer ring), with the locations of non-synonymous within gene variants indicated with black circles and the penetrance of each mutation in the population indicated by the size of the circle. The line graphs show the frequency of all variants (intergenic, synonymous, non-synonymous and nonsense) in each population. The vancomycin MIC for each population is also indicated by the shaded region. Mutations also identified in the respective end point clone (Fig. 1c) are highlighted by the coloured lines. A full list of all variants shown here is included in Supplementary Data 3.

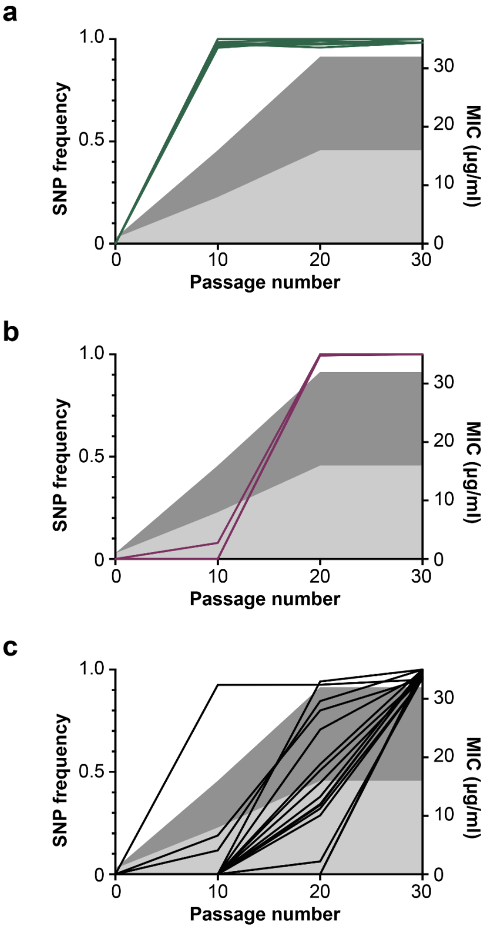


| Fixed at P10 | Fixed at P20 | Fixed at P30 |
|---|---|---|
| *CD0982* Bc9 | *CD0108* Bc4 | *CD0362* Bc11 |
| *CD1159* Bc4 | *CD0892* Bc10 | *CD0801* Bc5 |
| *CD3124* Bc2 | *CD0892* Bc10 | *CD0985* Bc7 |
| *CD3124* Bc7 | *CD0982* Bc10 | *CD1249* Bc7 |
| *CD3124* Bc9 | *CD0985* Bc8 | ***vanT* Bc3** |
| *CD3124* Bc10 | ***vanT* Bc4** | ***vanT* Bc11** |
| *CD3193* Bc2 | ***vanT* Bc5** | *CD1645* Bc11 |
| ***dacS Bc1*** | | *CD2081* Bc7 |
| ***dacS* Bc2** | | *CD2512* Bc9 |
| ***dacS* Bc9** | | *CD2579* Bc7 |
| ***dacR* Bc7** | | *CD2609* Bc11 |
| ***dacR* Bc10** | | *CD2614* Bc11 |
| | | *CD2649* Bc11 |
| | | *CD2852* Bc11 |
| | | *CD3141* Bc7 |
| | | ***dacJ* Bc9** |
| | | *CD3478* Bc11 |

**Supplementary Figure 5. Fixation of mutations during evolution.** Shown are the individual variants which fix (>95% penetrance) in all 10 parallel populations during vancomycin resistance evolution after 10 (**a**), 20 (**b**) and 30 passages (**c**). The frequency of each variant within their respective population is shown and the genes affected at each timepoint are shown in the table on the right. The genes in the *CD3437-9* cluster are highlighted in yellow and *vanT* (*CD1526*) in green. The range of vancomycin MICs observed across all populations (lowest, light grey; highest, dark grey) is indicated by the shaded areas.

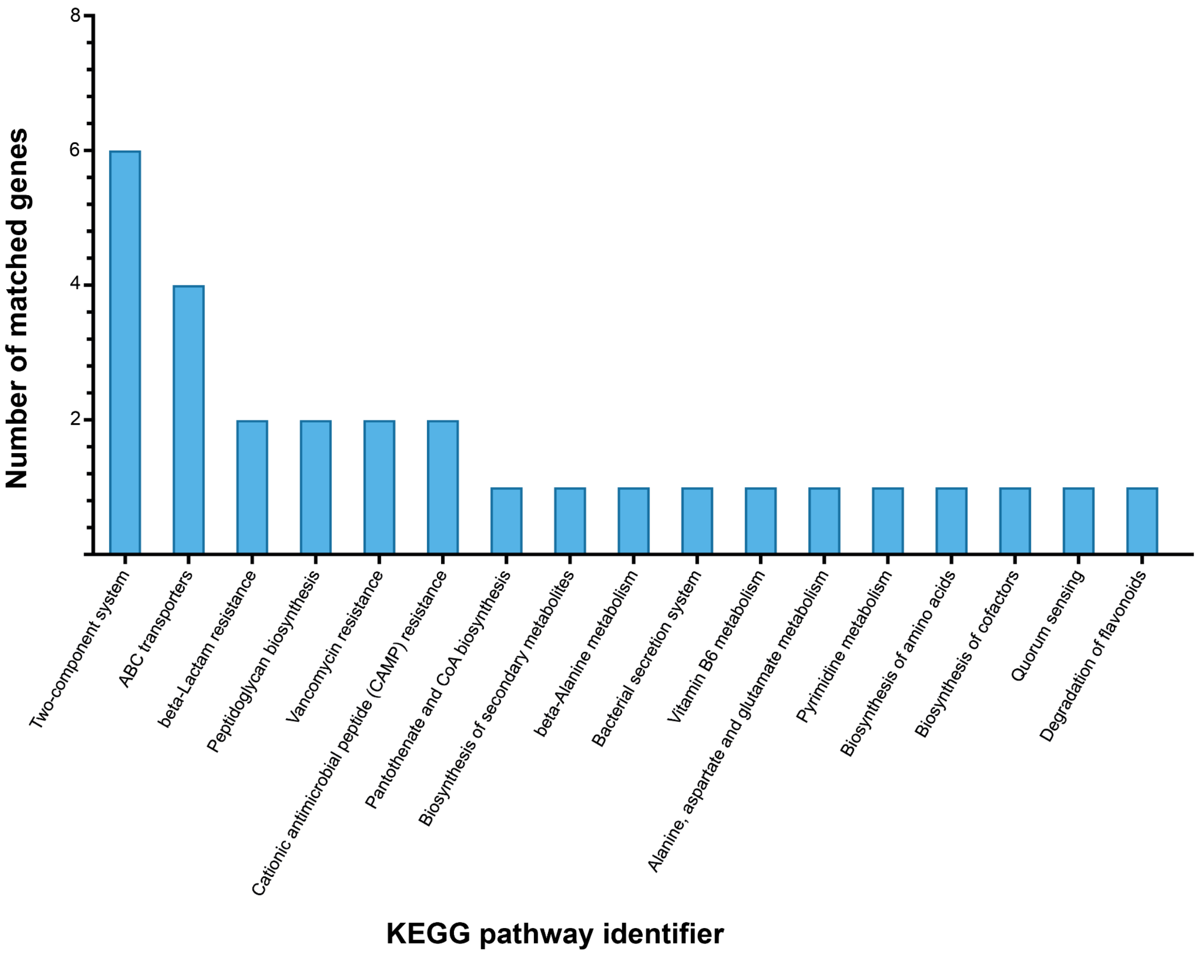


**Supplementary Figure 6. KEGG pathway analysis of genes identified in evolved populations.** Mutations occurring within genes (except those in the transiently hypermutating Bc1 P20) were coloured according to their barcoded replicate line, and pathways were visualised in KEGG (Kyoto Encyclopedia of Genes and Genomes) colour mapper[1]. Two-component systems and ABC transporters were the best-represented functional classes.

**Supplementary Table 1:** Strains used in this study

| Strain | Characteristics | Source |
|---|---|---|
| ***General Strains – C. difficile*** | | |
| R20291 | *C. difficile* ribotype 027 strain isolated during an outbreak at Stoke Mandeville hospital, UK in 2006. | [2] |
| R20291Δ*PaLoc* | R20291 with the entire pathogenicity locus (*tcdD, tcdB, tcdE, tcdA, dxtA)*, except the first codon of *dxtA*, deleted. | This study |
| R20291Δ*PaLoc*Δ*mutSL* | R20291Δ*PaLoc* with the entire *mutSL* locus (*mutS, mutL*), except the first codon of *mutS* and the last 2 codons of *mutL*, deleted. | This study |
| ***General Strains – E. coli*** | | |
| CA434 | *E. coli* conjugative donor. HB101 carrying R702. | [3] |
| NEB5α | *fhuA2 Δ(argF-lacZ)U169 phoA glnV44 Φ80Δ (lacZ)M15 gyrA96 recA1 relA1 endA1 thi-1 hsdR17.* | New England Biolabs |
| ***Barcoded Strains*** | | |
| R20291Δ*PaLoc pyrE::*barcode 1 | R20291Δ*PaLoc* with a 218 bp insertion between *CD0188* (*pyrE*) and *CD0189,* including 9 bp Barcode 1 (AAGTCCTCG) | This study |
| R20291Δ*PaLoc pyrE::*barcode 2 | R20291Δ*PaLoc* with a 218 bp insertion between *CD0188* (*pyrE*) and *CD0189,* including 9 bp Barcode 2 (TCTTGACCG) | This study |
| R20291Δ*PaLoc pyrE::*barcode 3 | R20291Δ*PaLoc* with a 218 bp insertion between *CD0188* (*pyrE*) and *CD0189,* including 9 bp Barcode 3 (AACAACACC) | This study |
| R20291Δ*PaLoc pyrE::*barcode 4 | R20291Δ*PaLoc* with a 218 bp insertion between *CD0188* (*pyrE*) and *CD0189,* including 9 bp Barcode 4 (AACAGGTGG) | This study |
| R20291Δ*PaLoc pyrE::*barcode 5 | R20291Δ*PaLoc* with a 218 bp insertion between *CD0188* (*pyrE*) and *CD0189,* including 9 bp Barcode 5 (ACCGATTAG) | This study |
| R20291Δ*PaLoc*Δ*mutSL pyrE::*barcode 7 | R20291Δ*PaLoc*Δ*mutSL* with a 218 bp insertion between *CD0188* (*pyrE*) and *CD0189,* including 9 bp Barcode 7 (CCTCCAACT) | This study |
| R20291Δ*PaLoc*Δ*mutSL pyrE::*barcode 8 | R20291Δ*PaLoc*Δ*mutSL* with a 218 bp insertion between *CD0188* (*pyrE*) and *CD0189,* including 9 bp Barcode 8 (CGAGGACAT) | This study |
| R20291Δ*PaLoc*Δ*mutSL pyrE::*barcode 9 | R20291Δ*PaLoc*Δ*mutSL* with a 218 bp insertion between *CD0188* (*pyrE*) and *CD0189,* including 9 bp Barcode 9 (CTGGTTCTA) | This study |
| R20291Δ*PaLoc*Δ*mutSL pyrE::*barcode 10 | R20291Δ*PaLoc*Δ*mutSL* with a 218 bp insertion between *CD0188* (*pyrE*) and *CD0189,* including 9 bp Barcode 10 (GGATGTTGG) | This study |

| | | |
|---|---|---|
| R20291Δ*PaLoc*Δ*mutSL* *pyrE::*barcode 11 | R20291Δ*PaLoc*Δ*mutSL* with a 218 bp insertion between *CD0188* (*pyrE*) and *CD0189,* including 9 bp Barcode 11 (GTCACCAGT) | This study |
| ***Evolved Strains*** | | |
| Bc1 | R20291Δ*PaLoc pyrE::*barcode 1 isolated after 60 days of vancomycin selection pressure. | This study |
| Bc2 | R20291Δ*PaLoc pyrE::*barcode 2 isolated after 60 days of vancomycin selection pressure. | This study |
| Bc3 | R20291Δ*PaLoc pyrE::*barcode 3 isolated after 60 days of vancomycin selection pressure. | This study |
| Bc4 | R20291Δ*PaLoc pyrE::*barcode 4 isolated after 60 days of vancomycin selection pressure. | This study |
| Bc5 | R20291Δ*PaLoc pyrE::*barcode 5 isolated after 60 days of vancomycin selection pressure. | This study |
| Bc7 | R20291Δ*PaLoc*Δ*mutSL pyrE::*barcode 7 isolated after 60 days of vancomycin selection pressure. | This study |
| Bc8 | R20291Δ*PaLoc*Δ*mutSL pyrE::*barcode 8 isolated after 60 days of vancomycin selection pressure. | This study |
| Bc9 | R20291Δ*PaLoc*Δ*mutSL pyrE::*barcode 9 isolated after 60 days of vancomycin selection pressure. | This study |
| Bc10 | R20291Δ*PaLoc*Δ*mutSL pyrE::*barcode 10 isolated after 60 days of vancomycin selection pressure. | This study |
| Bc11 | R20291Δ*PaLoc*Δ*mutSL pyrE::*barcode 11 isolated after 60 days of vancomycin selection pressure. | This study |
| ***Recapitulated strains*** | | |
| R20291Δ*PaLoc dacS*c.548T>C | R20291Δ*PaLoc* with *dacS* 548T>C point mutation identified in Evolved R20291Δ*PaLoc*Δ*mutSL* *pyrE::*barcodes 8 and 9 | This study |
| R20291Δ*PaLoc dacS*c.714G>T | R20291Δ*PaLoc* with *dacS* 714G>T point mutation identified in Evolved R20291Δ*PaLoc pyrE*::barcode 1 | This study |

**Supplementary Table 2:** Primers used in this study

| Oligonucleotide | Sequence | Use |
|---|---|---|
| ***Primers for Cloning*** | | |
| RF920 | cgtagaaatacggtgttttttgttaccctaTGGAATTTAGATATAAAAACCAATTC | Amplification of homology arm upstream of *PaLoc* with RF921 |
| RF921 | atttattttggtgtgGACAACATTGGAATTAAATCAG | Amplification of homology arm upstream of *PaLoc* with RF920 |
| RF922 | aattccaatgttgtcCACACCAAAATAAATGCC | Amplification of homology arm downstream of *PaLoc* with RF923 |
| RF923 | gggattttggtcatgagattatcaaaaaggCCCAACTATGGAAAAACC | Amplification of homology arm downstream of *PaLoc* with RF922 |
| RF2066 | AatacggtgttttttgttaccctagagctcCCACTTATAATTTCTAATGAAACTGTG | Amplification of homology arm upstream of *mutSL* with RF2067 |
| RF2067 | ccaaatattttacatCATTATCAAACCTCCTTCTTTTC | Amplification of homology arm upstream of *mutSL* with RF2066 |
| RF2068 | GGAGGTTTGATAATGATGTAAAATATTTGGATATTTAAAATATATGGAAAG | Amplification of homology arm downstream of *mutSL* with RF2069 |
| RF2069 | TTGGTCATGAGATTATCAAAAAGGGGATCCGCCCTTTAACTTGCACTC | Amplification of homology arm downstream of *mutSL* with RF2068 |
| ***Primers for Barcoding*** | | |
| RF1810 | GAAAAAGGCTTCTCTCATGAGAAG | To linearise pJAK081 to add barcode fragments |
| RF1811 | GGTACCATAAAAATAAGAAGCCTGC | To linearise pJAK081 to add barcode fragments |
| RF1902 | ACC GAAAAAGGCTTCTCTCATGAGAAG | Inverse PCR of pJAK201 to introduce barcode 3 |
| RF1903 | GTTGTT AAATGGAAGATGGAATAGAAGTAAGC | Inverse PCR of pJAK201 to introduce barcode 3 |
| RF1904 | GTGG GAAAAAGGCTTCTCTCATGAGAAG | Inverse PCR of pJAK201 to introduce barcode 4 |
| RF1905 | CTGTT AAATGGAAGATGGAATAGAAGTAAGC | Inverse PCR of pJAK201 to introduce barcode 4 |
| RF1906 | GATTAG GAAAAAGGCTTCTCTCATGAGAAG | Inverse PCR of pJAK201 to introduce barcode 5 |
| RF1907 | GGT AAATGGAAGATGGAATAGAAGTAAGC | Inverse PCR of pJAK201 to introduce barcode 5 |
| RF1912 | CAACT GAAAAAGGCTTCTCTCATGAGAAG | Inverse PCR of pJAK201 to introduce barcode 7 |
| RF1913 | GAGG AAATGGAAGATGGAATAGAAGTAAGC | Inverse PCR of pJAK201 to introduce barcode 7 |

| | | |
|---|---|---|
| RF1914 | GACAT GAAAAAGGCTTCTCTCATGAGAAG | Inverse PCR of pJAK201 to introduce barcode 8 |
| RF1915 | CTCG AAATGGAAGATGGAATAGAAGTAAGC | Inverse PCR of pJAK201 to introduce barcode 8 |
| RF1916 | GTTCTA GAAAAAGGCTTCTCTCATGAGAAG | Inverse PCR of pJAK201 to introduce barcode 9 |
| RF1917 | CAG AAATGGAAGATGGAATAGAAGTAAGC | Inverse PCR of pJAK201 to introduce barcode 9 |
| RF1918 | TTGG GAAAAAGGCTTCTCTCATGAGAAG | Inverse PCR of pJAK201 to introduce barcode 10 |
| RF1919 | CATCC AAATGGAAGATGGAATAGAAGTAAGC | Inverse PCR of pJAK201 to introduce barcode 10 |
| RF1920 | CAGT GAAAAAGGCTTCTCTCATGAGAAG | Inverse PCR of pJAK201 to introduce barcode 11 |
| RF1921 | GTGAC AAATGGAAGATGGAATAGAAGTAAGC | Inverse PCR of pJAK201 to introduce barcode 11 |
| ***Primers for qPCR*** | | |
| RF2504 | CATCATTACCAGGTGTAGCAGTG | Amplification of ~200bp *rpoA* fragment for qPCR |
| RF2505 | GGAGGACAGATTATATCTGCACC | Amplification of ~200bp *rpoA* fragment for qPCR |
| RF2506 | CAATCACATCATTAGCAATTTATTCCATG | Amplification of ~200bp *dacS* fragment for qPCR |
| RF2507 | GTTCATCAATATCATCCTTTTCTTTATCC | Amplification of ~200bp *dacS* fragment for qPCR |
| RF2508 | GGATGGGATAGAAGTTTGTAGAAAAG | Amplification of ~200bp *dacR* fragment for qPCR |
| RF2509 | CTCTTCTAATCAGTGATTTCACTCTC | Amplification of ~200bp *dacR* fragment for qPCR |
| RF2510 | CAACATGATTCAGAACAAGATGTTGAG | Amplification of ~200bp *dacJ* fragment for qPCR |
| RF2511 | GCTTGCTTAACTAAATCTTCAACTGC | Amplification of ~200bp *dacJ* fragment for qPCR |
| RF2512 | CTGATTTTAGAAAAGTATATAAACACGG C | Amplification of ~200bp *rnpA* fragment for qPCR |
| RF2513 | CTTGCTATAAATACTATATCATATCCAGG | Amplification of ~200bp *rnpA* fragment for qPCR |

**Supplementary Table 3:** Plasmids used in this study

| Plasmid | Characteristics | Source |
|---|---|---|
| ***General Plasmids*** | | |
| pJAK112 | pMTL-SC7215 based vector with added BamHI and SacI restriction sites for cloning. | [4] |
| pJAK143 | *PaLoc* deletion – 1200bp homology arms upstream and downstream of the *PaLoc* for deletion of the entire pathogenicity locus (*tcdD, tcdB, tcdE, tcdA, dxtA)*, except the first codon of *dxtA*. | [5] |
| pJEB002 | *mutSL* deletion – 1200 bp homology arms upstream and downstream of *mutSL* for deletion of the entire *mutSL* locus (*mutS, mutL*), except the first codon of *mutS* and the last 2 codons of *mutL.* | This study |
| pMTL-SC7215 | Allele exchange vector for *codA*-based selection. | [6] |
| ***Plasmids for Barcoding*** | | |
| pJAK081 | pMTL-SC7215 based vector with 1,200 bp homology arms for insertion of DNA sequences between *CD0188* (*pyrE*) and *CD0189* in the *C. difficile* R20291 genome. | This study |
| pJAK201 | *pyrE::*barcode 1 – pJAK081 based vector, with a 218 bp insertion containing 9 nt Barcode 1 (AAGTCCTCG) | This study |
| pJAK202 | *pyrE::*barcode 2 - pJAK081 based vector, with a 218 bp insertion containing 9 nt Barcode 2 (TCTTGACCG) | This study |
| pJAK203 | *pyrE::*barcode 3 - pJAK081 based vector, with a 218 bp insertion containing 9 nt Barcode (AACAACACC) | This study |
| pJAK204 | *pyrE::*barcode 4 - pJAK081 based vector, with a 218 bp insertion containing 9 nt Barcode 4 (AACAGGTGG) | This study |
| pJAK205 | *pyrE::*barcode 5 - pJAK081 based vector, with a 218 bp insertion containing 9 nt Barcode 5 (ACCGATTAG) | This study |
| pJAK207 | *pyrE::*barcode 7 - pJAK081 based vector, with a 218 bp insertion containing 9 nt Barcode 7 (CCTCCAACT) | This study |
| pJAK208 | *pyrE::*barcode 8 - pJAK081 based vector, with a 218 bp insertion containing 9 nt Barcode 8 (CGAGGACAT) | This study |
| pJAK209 | *pyrE::*barcode 9 pJAK081 based vector, with a 218 bp insertion containing 9 nt Barcode 9 (CTGGTTCTA) | This study |
| pJAK210 | *pyrE::*barcode 10 - pJAK081 based vector, with a 218 bp insertion containing 9 nt Barcode 10 (GGATGTTGG) | This study |
| pJAK211 | *pyrE::*barcode 11 - pJAK081 based vector, with a 218 bp insertion containing 9 nt Barcode e 11 (GTCACCAGT) | This study |
| ***Plasmids for Recapitulating*** | | |
| pJEB019 | *dacS*c.548T>C - pJAK112 based vector containing 1,926 bp homology arms centred on a *dacS* 548T>C point mutation. | This study |
| pJEB026 | *dacS*c.714G>T - pJAK112 based vector containing 1,926 bp homology arms centred on a *dacS* 714G>T point mutation. | This study |
| ***qRT-PCR Plasmids*** | | |

| | | |
|---|---|---|
| pJEB029 | qPCR 1 - pUC-GW-Kan vector including ~200 bp fragments of *rpoA, dacS, dacR, dacJ* and *rnpA* for qRT-PCR | This study |

**Supplementary Table 4**. DNA Accession numbers
